# General calibration of microbial growth in microplate readers

**DOI:** 10.1101/061861

**Authors:** Keiran Stevenson, Alexander F. McVey, Ivan B.N. Clark, Peter S. Swain, Teuta Pilizota

## Abstract

Optical density (OD) measurements of microbial growth are one of the most common techniques used in microbiology, with applications ranging from antibiotic efficacy studies, studies of growth under different nutritional or stress environments, to studies of different mutant strains, including those harbouring synthetic circuits. OD measurements are performed under the assumption that the OD value obtained is proportional to the cell number, i.e. the concentration of the sample. However, the assumption holds true in a limited range of conditions and calibration techniques that determine that range are currently missing. Here we present a set of calibration procedures and considerations that are necessary to successfully estimate the cell concentration from OD measurements.

## Introduction

Bacteria and yeast are widely studied microorganisms of great economic, medical and societal interest. Much of our understanding of bacterial and yeast life cycles stems from monitoring their proliferation in time and the most routine way of doing so is using optical density (OD) measurements. The applications range from routine checks during different cloning techniques;^1^ through studying cellular physiology and metabolism;^2^,^3^to determining the growth rate for antibiotic dosage^4^,^5^and monitoring of biomass accumulation during bio-industrial fermentation.^6^ Here we introduce a set of calibration techniques that take into account the relevant parameters affecting OD measurements, including at high culture densities, in a range of conditions commonly used by researchers. OD measurements have become synonymous with measurements of bacterial number (*N*) or concentration (*C*), in accordance with the Beer-Lambert law. However, OD measurements are turbidity measurements,^7^,^8^ thus, the Beer-Lambert law can be applied, with some considerations, only for microbial cultures of low densities. OD measurements in plate readers, increasingly used for high-throughput estimates of microbial growth, operate predominantly at higher culture densities where OD is expected to have a parabolic dependency on *N*.^8^ Additionally, the proportionality constants (either in low or high density regimes) strongly depend on several parameters, for example cell size, which need to be considered and included in robust calibration techniques. Yet,these techniques, essential when using OD measurements for quantitative studies of microbial growth,including growth rates, lag times and cell yields, have thus far not been established.

The Beer-Lambert law (Supplementary Note 1) assumes *OD*∼*C*, which is true if the light received by the detector of a typical spectrometer is the light that did not interact with the sample in any way.^7^,^9^ In general, when microbial cells are well dispersed in the solution (for the cases where *N* is small,i.e. single scattering regime) and the geometry ofthe spectrometer is suitable, the Beer-Lambert law is a good approximation for turbidity measurements and *N* (or *C*) is ∼ *OD*.^7^, ^8^The suitable geometry of the spectrometer refers to the likelihood of the scattered light reaching the detector in the spectrometer even in the single scattering regime (Fig. 1A-B).Most bacteria and yeast scatter light at small angles (a few degrees^10^). Thus, the distance from the scatterer to the detector (*d*) and the radius (*R*) of the aperture (Fig. 1A) will determine how close a particular spectrometer is to the ideal case, even in the single scattering regime. Supplementary Figure 1A shows measurements takenofthe same sample in five different spectrometers, indicating that different spectrometers need to be cross-calibrated even when used in the single scattering regime.^9, 11^ As *N* increases the probability of incident light being scattered by particles multiple times also increases (Fig. 1B), the so called multiple scattering regime. In this regime the Beer-Lambert law is no longer a suitable approximation andOD is expected to have a parabolic dependency on *N*.^8^ Similarly, as the probability of multiple scattering events increases even further (i.e. for very high *N*), light is increasingly deflected away from the detector, and can be described with diffusive approximations (so called photon diffusion limit)^12, 13^.

**Fig. 1.**
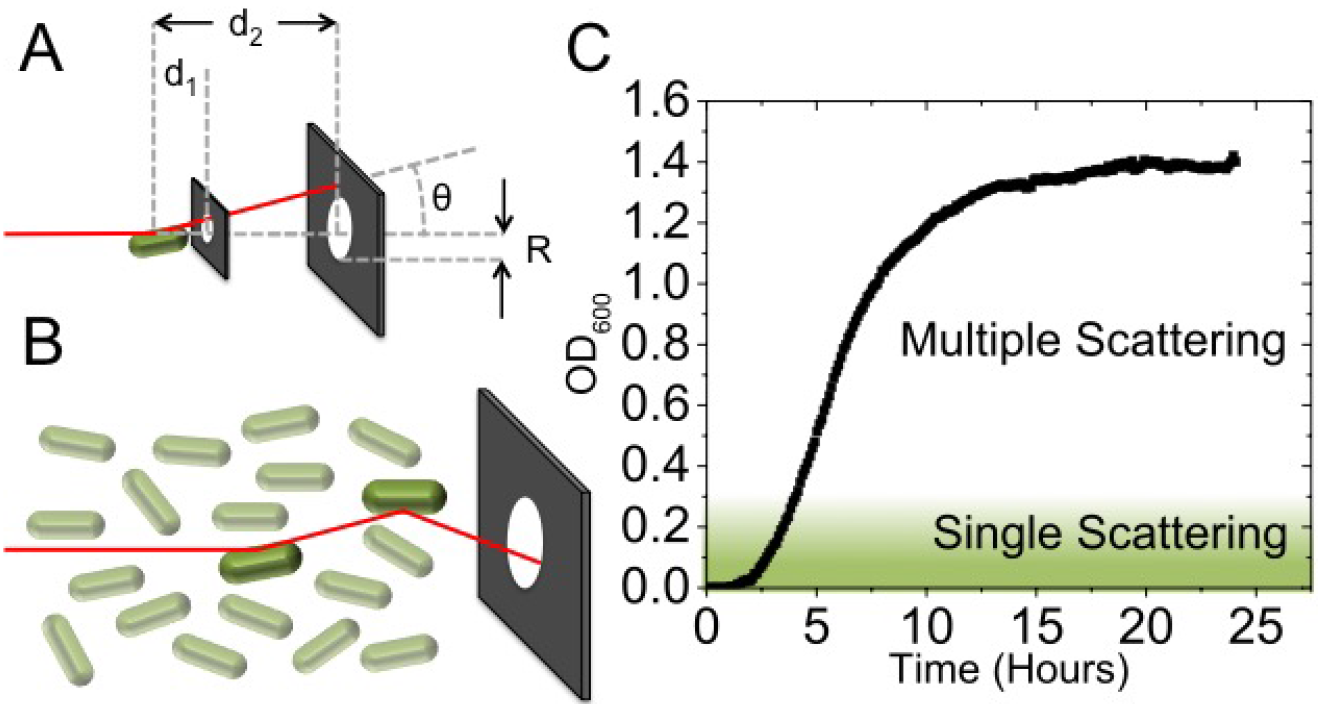
Schematic showing that light incident on a sample is scattered by an angle θ from the optical axis (*z*), either once as in (A) or multiple times (B).Single scattering events are more likely to deflect light away from the aperture (radius *R*), but the effect of this scattering is highly dependent upon the size of the detector and the distance between detector and sample (*d_1_* or *d_2_*). As the concentration of cells increases, the probability of light being scattered back into the detector is increased. (C) Atypical OD curve for *E. coli* measured at λ =600 nm (for measurements at different λsee Supplementary Fig. 2). Single and multiple scattering regimes have been indicated with different colour shading. The OD saturates when *N* is large enough to deplete the nutrient sources in the media

Figure 1C shows a typical OD curve for *Escherichia coli* grown in rich media, with single and multiple scattering regimes indicated. In the single scattering regime (small *N* and OD_600_≲0.2^8,13,14^) that can be approximated as the Beer-Lambert law (*OD* ∼ *N*), exact solutions to the scattering problem exist.^9^ Depending on the size and shape of the scatterer (bacteria or yeast), as well as the index of refraction difference between the scatterer and the media, different approximations have been used to solve the single scattering regime problem (Supplementary Table 1). For example, the Jobst approximation, used for spherical bacteria with dimensions comparable to the wavelength of light (*r* ∼ λ, where *r* is radius of the bacteria and λ the wavelength of incident light), gives OD to be proportional to *N* and *r*^4^ (i.e. bacterial volume to the power of four thirds).^15^ Multiple scattering effects can be incorporated into scattering theory with the inclusion of a correction factor *CF*(σ,*z*) (Supplementary Note 2). Unfortunately, in order to calculate *CF*, *N* must be known. Therefore, when using microplate readers in the multiple scattering regime the most practical way of determining the relationship between OD and *N* is to calibrate. Correct calibration is of particular relevance to high-throughput quantitative growth rate and cell yield measurements in platereaders and bio-reactors, but it is rarely reported by researchers. Additionally, it is often assumed that a single calibration curve for a given instrument is sufficient,^16^ or calibration is performed by counting colony forming units (CFUs),^17^ which only takes into account live bacteria and is not necessarily suitable for growth under different antibiotics (where cells can be dead, but not lysed). For correct calibration, shape and size changes of scatterers (bacteria and yeast) as well as index of refraction changes of the media (*n_m_*) and/or scatterers (*n_p_*) need to be taken into account.

In order to demonstrate the effect of differences in the size of scatterers on OD in single and multiple scattering regime, we measured monodisperse solutions of beads with different diameters (*D*), known index of refraction and known concentrations (*C*) determined by direct counting in a microscope. Calibration curves for each sample of beads are given in Figure 2A. For a fixed OD, *C* increases as the diameter of the scatter decreases (Fig. 2B). The relative effect of *C* on OD is more pronounced for *D* ∼ λ, in both single and multiple scattering regimes,indicating that for yeast cultures (with typical cell size ∼5 μm) the changes in size should affect the calibration curve less than for bacterial cultures (with typical cell size on the order of ∼1 μm). Similar conclusions holds true for samples heterogeneous in size. Upon introduction of a polydisperse sample of 0.5 μm and 1.0 μm diameter beads (a 1:1 mixture by volume) *C* is seen to deviate greatly from both monodisperse solutions (Fig. 2C). Consequently, while the OD versus *C* calibration curves follow similar trends (a second order polynomial^8^), the exact calibration curve is highly dependent upon *D*, particularly for *D* ∼ λ.

The effects of changing the size of the scatterer seen in bead suspensions are also visible in cultures of *E.coli* and yeast measured in a platereader (Fig. 3A and Methods). The representative images (Fig. 3B-H) show the change in cell geometry. Different cell sizes for *E.coli* were obtained by sampling at different stages of growth in rich undefined media (Supplementary Fig. S4 and by growing the culture in the presence of a sublethal concentration of ampicillin, a β-lactam antibiotic that inhibits the formation of thecell wall (Fig. 3B). Ampicillin induces filamentation^18,19^(Fig. 3E), a property shared by many other stresses(including antibiotic groups cephalosporins and quinolones^20^ and UV irradiation and oxidative damage^21^). To obtain different sizes of yeast cells we used three different strains (wild type haploid and diploid, and a diploid mutant exhibiting increased cell size).^22^ Each chosen scatterer in Figure 3 exhibits a different relationships between OD and *C*. In particular, thecalibration curve obtained using filamentous cells is significantly altered whencompared tothe rest of the *E.coli* samples. The differences in the calibration of yeast samples are reduced when compared to bacterial cultures, with a small change in the calibration as *D* increases from 4μm to 6μm.

**Fig. 2.**
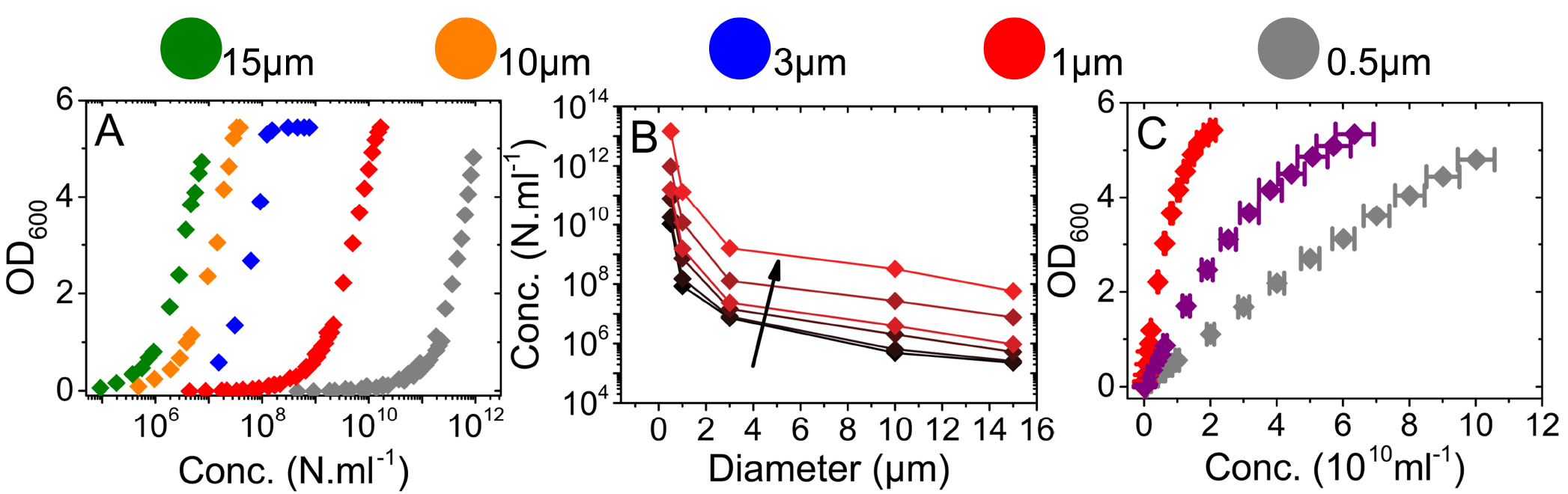
OD measurements of spherical polystyrene bead suspensions. Each set of data consists of dilutions of a single stock whose actual concentration was determined by counting ina microfluidic slide (see Methods). Bead size corresponds to the diagram above the graphs (0:51±0:01 μm, 0:96±0:07 μm, 3:00±0:07 μm, 10:0±0:6 μm and 15:7±1:4 μm) and bead index of refraction is *n_p_* = 1:59. Representative images of beads used for *C* measurements are shown in Supplementary Figure 3. (A)Comparison of concentrations (see Methods) and OD measured i a microplatereader for a given bead diameter. (B) The bead concentration as a function of *D* obtained from(A) for the following ODs: 0.05, 0.1, 0.5, 1 and 10. Increasing OD is represented as increasing brightness of red and by the arrow. (C) Measurements of 0.5 μm and 1:0 μm bead suspensions and the resultant (1:1 by volume) mixture in purple.Figure

**Fig. 3.**
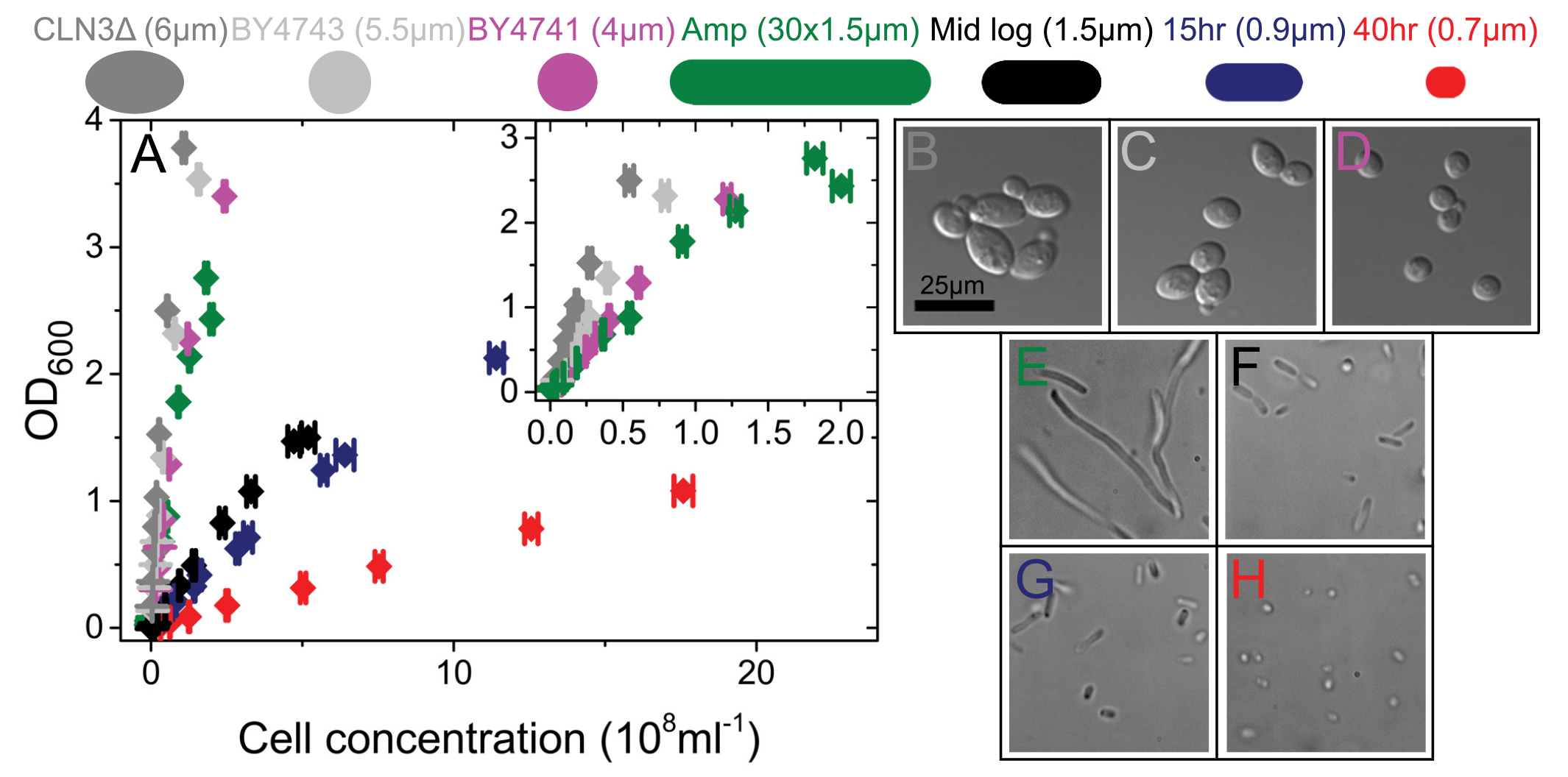
(A) OD vs *C* is shown for yeast-diploids (grey);-haploid (purple); filamentous *E. coli* (green) and mid-log (black);early (blue) and late (red) stationary phase *E. coli* using dilutions of a single stock whose actual cell concentrations were determined by counting in a microscope. (B-H) Representative images of each culture obtained during microscopy. Scale bar is shown in B and applied to all panels. B: Yeast CLN3Δ homozygous diploid mutant C: Yeast diploid, D: Yeast haploid. E: *E. coli* cells grown in the presence of sub lethal concentration of ampicillin to induce filamentation. F: *E. coli* cells grown to mid log phase in LB (OD= 0:2). G: *E. coli* cells at early stationary phase (OD = 2:3),H: *E. coli* cells at late stationary phase (After40 hr)

**Fig. 4.**
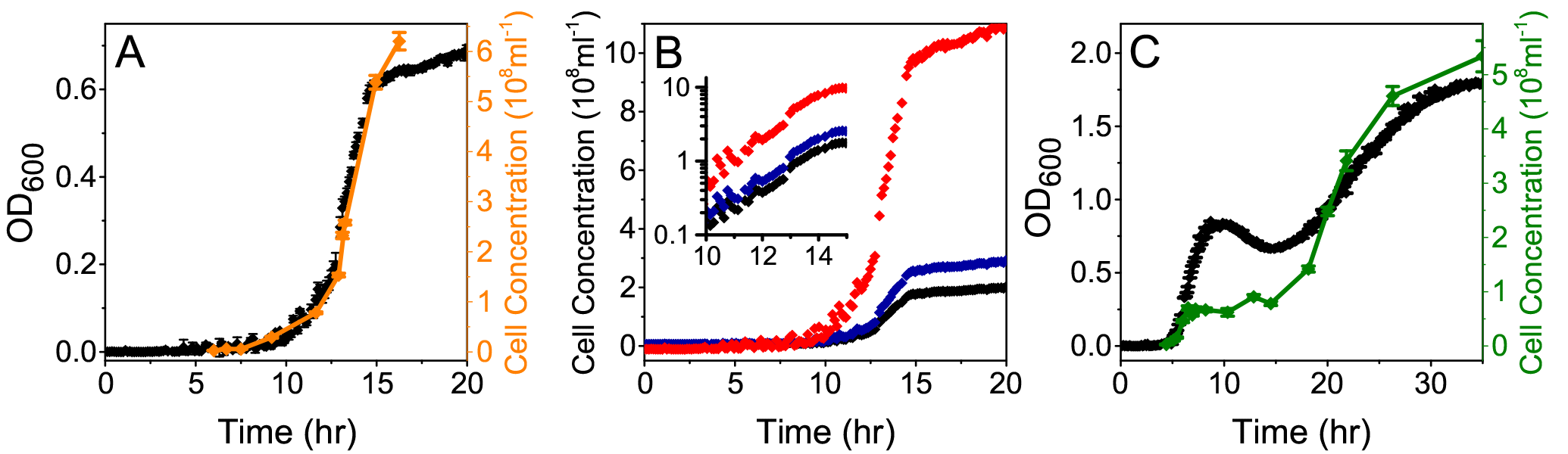
Growth curves and cell counts. (A) OD (black) and cell concentration (orange) during growth in MM9 with glucose. OD and *C* are closely correlated until starvation when cell size is no longer constant. (B) The OD curve from (A),converted to cell concentration using the the mid log (black), 15 hr (blue) and 40 hr (red) curves in Figure 3A. Calibration performed on cells of different sizes produce large differences in cell concentration. (C) OD (black) and cell concentration(green) in LB with 9 μg ml^−1^ Ampicillin. Poor correlation between OD and *C* is caused by antibiotic induced filamentation and fluctuating filament lengthduring growth.

The effects of changes in the calibration curve during a single growth curve are demonstrated in Figure 4 and Supplementary Figure S5 by counting the cell concentration in parallel with the measurement of the OD. We first demonstrate that if the cell size does not change during culture growth, the calibration curve maintains parabolic dependency on *N*. In order to keep a constant cell size we grew *E*. *coli* in MM9 with glucose as the sole carbon source (Methods). Under these conditions OD and *C* are well correlated throughout the exponential growth, only diverging at 15 hr when the carbon source is depleted and the cells enter stationary phase, therefore reducing in volume (Fig. 4A). However, if cells are changing size during growth, for example when growing on rich undefined media (Supplementary Fig. 5), the calibration curve will change in time as well.To demonstrate the effect of using calibration curves obtained for cells of different sizes,in Figure 4B we show the OD curve from Figure 4A converted to *C* using three different calibrations curves. Conversion using cells taken from mid log phase of growth (black, also black in Fig. 3A), after 15 hr (blue) and 40 hr (red) is shown. *C* is significantly altered depending on the calibration curve selected, with the effect on the lag time and final cell yield particularly pronounced.

To further demonstrate the effect on changes in cell size throughout the growth curve we grew *E.coli* under sublethal concentrations of ampicillin (Fig. 4C). Significant deviation between OD and *C* is visible. During the initial part of the log phase OD and *C* show the same time dependency. However, at OD ∼ 0.2, *N* remains roughly constant while the OD increases. The increase in OD for constant *N* is the result of an increase in cell mass rather than increasing *N* (Fig. 3E). The decrease in OD at constant *N* (10 hr) is most likely the result of division of filaments into smaller cells. 15 hr after culture inoculation, both *N* and OD increase again as filamentous bacteria both divide and grow as smaller cells.

To demonstrate the difference between the expected OD and *C* relationship for scatterers of a fixed size and those whose size is changing during growth, in Supplementary Figure 5 we show comparison of OD and *C* from Figure 4A and Figure 4C. In the case ofa constant cell size (Supplementary Fig. 5A) the calibration curve follows the same second degree polynomial expected from Figure 3A and Figure 2A and Figure 2C, whereas for growth in LB+ampicillin, where cell size changes, the calibration curve is more complex (Supplementary Fig. 5B). Supplementary Figure 6 shows the difference between growth rates obtained from non-calibrated OD and *C* measurements for the same cell culture. The value and time point at which maximum growth rate is reached in Supplementary Figure 6A (corresponding to Fig. 4A) are similar, whereas Supplementary Figure 6B shows that maximum growth rate obtained from *C* is reached several hours before the maximum growth rate obtained from OD measurements. Additionally, Supplementary Figure 6B shows that the two values differ significantly, by a factor of two.

Apart from size of the scatterer, changes in the difference between refractive index of the growth media (*n_m_*) and the refractive index of the scatterer (*n_p_*) can have a significant effect on the OD calibration curve. To demonstrate this effect we changed *n_m_* by the addition of sucrose, while keeping the refractive index of the scatterer (1 μm bead) the same (Supplementary Fig. 7). As the relationship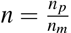decreases the OD of a fixed *N* is similarly reduced. The effect is small for beads as the relative difference between *n_p_* and *n_m_* is large, but will be more pronounced for biological samples like bacteria, as *n* of bacteria is smaller (Supplementary Table 2).

To investigate the effect of bacterial lysis and intracellular matter leaking into the media on *n_m_*, for example during growth under antibiotics, we measured the refractive index of LB media with different concentrations of lysed *E*. *coli* cells. The intracellular material (Supplementary Table 3 and Supplementary Fig. 8), released into the media from high concentrations of cells (as high as 2 · 10^9^ml^−1^ cells)results in only a small increase in *n_m_*. The increase is significantly lower than *n_m_* variations caused by the introduction of even low concentrations of sucrose to ddH_2_O (Supplementary Fig. 8). Thus, cell lysis as a result of growth under different antibiotics will unlikely change *n_m_* sufficiently to alter the OD versus *C* calibration curve. However, growth at high sugar concentrations, such as for food industry applications, will.

We have presented calibration considerations and protocols needed for quantitative measurements of microbial growth rates based on OD measurements. We show that different spectrometers and microplate readers need to be cross-calibrated to compare the OD readings as an absolute number. Furthermore, variations in *D* and refractive index of the cell or the media need to be considered and calibrated in order to avoid significantly over-or underestimating the number of cells present in the sample. Thus, we recommend first determining if significant changes in cell size are expected during growth of microbial culture. If this is not the case, and size is expected to remain constant, calibration of OD against *N* needs to be performed once for each *D* and index of refraction, and ideally reported in publications. The closer the *D* of the scatterer to λ the more important it is to perform calibration of OD against *N* for each different cell size. Changes in refractive index can beparticularly relevant duringgrowth in media with high sugar concentrations (such as those in food sciences^26^ and drinks with high osmolarity (such as beer)). We have shown that changes in *n_m_* due to lysis induced leakage of cell material, for example when grown in the presence of antibiotics, are small. However, we note that the effect of changing media index of refraction is likely to be more pronounced in bioreactor experiments where *C* is far larger. If cell size is expected to change significantly during the course of growth of the microbial culture (for example: growth under antibiotics or various other stresses,growth of shape inducing mutants, growth of overexpression strains, growth of strainsthat nduce chain or clump formations) OD measurements are no longer suitable and direct counting of *N* should be performed, using, for example, microscopy.

## Methods

### Bacteria cultures

All bacterial cell culture studies were conducted using *E.coli* BW25113 (F^−^, DE(araD-araB)567, lacZ4787(del)::rrnB-3, LAM^−^, rph-1,DE(rhaD-rhaB)568, hsdR514), a close relative of MG1655, with plasmid pWR20 which expresses and enhanced GFP for cytoplasmic volume monitoring.^27^ All experiments were conducted in LB media except where explicitly stated. MM9 medium contained 0.1 % Glucose, noaminoacids and 20 mmol KCl. MM9 (Modified M9) is of the same composition as M9^28^ exceptsodium phosphate buffer only was used. Where cells were grown with antibiotics, 9 μgml^−1^ Ampicillin was added to the culture medium before the addition of cells.

### Yeast cultures

Yeast studies used three S288c-derived strains: BY4741, BY4743^29^ and the cln3 homozygous deletion derived from the *Saccharomyces* genome deletion project.^30^ Cells were cultured at 30 °C in YPD media, containing 2% glucose.

### Colloidal bead cultures

Colloidal bead cultures were created using dilutions of polystyrene beads of known diameter (*D*) 0.51±0.01 μm (Polysciences), 0.96±0.07μm(Bangs Laboratory), 3.00±0.07μm, 10.0±0.6μm and 15.7±1.4μm (all Polysciences) and known index of refraction (*n_p_*=1.59) in ddH_2_0. At each *D*, a dilution series was performed,to produce samples of known concentrations (*C*) between 10^5^ and 10^12^ Nml^−1^. For all samples *C* was experimentally determined by counting the number of beads in 10 μl of the dilution in a microscope tunnel slide.^27^

### Optical density measurements

OD measurements of bacterial and colloidal bead cultures were performed in a Spectrostar Omega microplate reader (BMG, Germany) with a Costar Flat Bottom 96-well plate with lid and 200 μl per well (300 μl for data in Fig. 4). Absorbance was measured at wavelength 600nm and temperature 37 °C and the mean of 5 readings taken. For bacterial cultures, 30 wells were grown to OD=0.15 in MM9 medium. The wells were pooled and 125 μg ml^−1^ chloramphenicol was added to inhibit further cell division or growth. The cells were then diluted in increments to provide a range of OD readings. A single dilution of cells for each series was then imaged in the brightfield microscope as above for the polystyrene beads. All measurements in the main text were reported using the BMG with correction values, which is given as the measured OD multiplied by 1.0560 for 300 μl, 1.5848 for 200 μl and 6.3694 for 100 μl.

OD measurements for yeast cultures were performed in a Tecan M200 fluorescent spectrometer, using a Costar Flat Bottom 96-well plate with lid and 200 μl per well for all measurements. Cells were cultured for 16 hr in YPD media and dilutions for measurement made in the same media. Duplicate OD measurements were taken at 600 nm (bandwidth 9) with 15 flashes at 30 °C. Yeast cell counts were performed using a Neubauer improved bright-line haemocytometer (Marienfeld).

Calibration between the two spectrometers was performed using *E. coli* grown in LB to mid log as in Figure 3A black and the OD measured in both platereaders. The relative difference in measurements was then calculated and used to correct the data gathered for yeast. The calibration is shown in Supplementary Figure 1B.

### Brightfield Microscopy

Imaging of samples was performed using a custom-built brightfield microscope consisting of a 100× oil immersion objective lens (Nikon) with the sample mounted on a Nano-LP200piezoelectric stage (Mad City Labs). Illumination of the sample was provided by a white LED (Luxeon Star) and images recorded on an iXon Ultra 897 EMCCD camera (Andor). Stacks of images through each sample were acquired every 0.05 s and separations of 1 μm, ensuring all scatterers in the volume were identified without introducing overcounting. True values of *C* were experimentally determined by counting *N* present in the known stack volume determined by the field of view of the microscope (55.6 × 55.6× 100μm).

### Osmolarity measurements

For osmolarity measurements, beads (*D* =1 μm) were diluted from manufacturer stock solution into ddH_2_O and then further diluted into solutions of 0mOsm, 116mOsm, 231 mOsm, 463 mOsm and 925 mOsm (achieved by diluting sucrose (Sigma) in ddH_2_O) to produce concentrations in media of refractive index *n_m_*=1.333;1.339;1.344;1.353 and 1.368 respectively. For each osmolarity,*C* was determined by counting *N* using brightfield microscopy as above.

### Fitting of bead optical density measurements

Fitting of the data presented in Figure 2A was performed in the Matlab^31^ environment utilising the built in curve fitting tools. Data was first trimmed to remove points where the spectrometer had saturated then fitted as a 2^nd^ degree polynomial with robust fitting using the bisquare method. The polynomials found are presented in Supplementary Table 4. These polynomials were then solved for OD=0.05,0.1,0.5,1 and 10 to give the traces presented in Figure 2C.

### *E. coli* cell concentration monitoring during growth curves

In order to determine *C* during batch culture growth, cells were added to 10 ml of the experimental media, which was then divided among 30-60 wells of a 96-well plate. The cells were grown at 37 °C and OD measured every 7.5 min. When an increase in OD above the baseline was observed sampling for *C* began, with each new sample taken from a separate well. For all growth curves at least 8 wells were left untouched to provide a complete growth curve for comparison. In the rare cases where the growth curve deviated significantly from the average, those traces were excluded from all measurements. 125 μg ml^−1^ chloramphenicol was added to samples taken from the plate to arrest all cell division and growth. Samples were then imaged under brightfield illumination and *N* counted manually to determine *C* for each time point. At OD>0.2 samples were diluted in the culture medium to reduce cell overlap in the slide. For Figure 4 the two data sets were aligned by using a least squares method to determine the scales of both axes, minimising the sum of: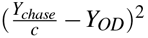 where *c* is the scaling factor. For Supplementary Figure 5 solid and dashed fits were produced using 1^st^ and 2^nd^ degree polynomials respectively. The fitted region for each was selected by expanding sequentially from zero until the fit quality started to drop.

### Refractive index measurements

*E. coli* cells (strain MG1655) were grown in LB to OD=0.3, spun down and concentrated 33.3 × before being subjected to sonication to lyse the cells. This cell lysate was diluted to 0.0625×, 0.125×, 0.25×, 0.5× and 1× concentrations and the refractive index measured. The original cell extract was counted as above to determine the concentration of cells before lysing. DNA was obtained as 23-mer primers (Sigma-Aldrich) suspended in ddH_2_O, with concentration measured usinga NanoDrop (ThermoScientific). Refractive indices of solutions were measured in a manual refractometer (Bellingham and Stanley, London). Sucrose data was obtained from a standard brix index.^32^

## Acknowledgements

KS is supported by the BBSRC iCASE grant to TP. AFM, IBNC, PSS and TP acknowledge the support of UK Research Councils Synthetic Biology for Growth programme and KS, AFM, IBNC, PSS and TP are members of a BBSRC/EPSRC/MRC funded Synthetic Biology Research Centre. TP acknowledges the support of BBSRC CBMNet grant. We thank Jerko Rosko, Dejan Kunovac and all other members of the Pilizota and Swain lab for their comments and support, Andrew Schofield for granting as access to the refractometer and Meriem El Karoui and Rosalind Allen for helpful comments and discussions.

### Author Contributions

TP and KS designed the study. TP, KS and PSS conceived the experiments. TP, KS and IBNC performed the experiments and analysed the data. KS, AFM and TP interpreted the data and wrote the paper. All authors reviewed the manuscript.

### Competing Financial Interest

The authors declare no competing financial interests.

